# Distinct mutations and lineages of SARS-CoV-2 virus in the early phase of COVID-19 pandemic and subsequent one-year global expansion

**DOI:** 10.1101/2021.01.05.425339

**Authors:** Yan Chen, Shiyong Li, Wei Wu, Shuaipeng Geng, Mao Mao

## Abstract

A novel coronavirus, SARS-CoV-2, has caused over 190 million cases and over 4 million deaths worldwide since it occurred in December 2019 in Wuhan, China. Here we conceptualized the temporospatial evolutionary and expansion dynamics of SARS-CoV-2 by taking a series of cross-sectional view of viral genomes from early outbreak in January 2020 in Wuhan to early phase of global ignition in early April, and finally to the subsequent global expansion by late December 2020. Based on the phylogenetic analysis of the early patients in Wuhan, Wuhan/WH04/2020 is supposed to be a more appropriate reference genome of SARS-CoV-2, instead of the first sequenced genome Wuhan-Hu-1. By scrutinizing the cases from the very early outbreak, we found a viral genotype from the Seafood Market in Wuhan featured with two concurrent mutations (i.e. M type) had become the overwhelmingly dominant genotype (95.3%) of the pandemic one year later. By analyzing 4,013 SARS-CoV-2 genomes from different continents by early April, we were able to interrogate the viral genomic composition dynamics of initial phase of global ignition over a timespan of 14-week. 11 major viral genotypes with unique geographic distributions were also identified. WE1 type, a descendant of M and predominantly witnessed in western Europe, consisted a half of all the cases (50.2%) at the time. The mutations of major genotypes at the same hierarchical level were mutually exclusive, which implying that various genotypes bearing the specific mutations were propagated during human-to-human transmission, not by accumulating hot-spot mutations during the replication of individual viral genomes. As the pandemic was unfolding, we also used the same approach to analyze 261,323 SARS-CoV-2 genomes from the world since the outbreak in Wuhan (i.e. including all the publicly available viral genomes) in order to recapitulate our findings over one-year timespan. By 25 December 2020, 95.3% of global cases were M type and 93.0% of M-type cases were WE1. In fact, at present all the four variants of concern (VOC) are the descendants of WE1 type. This study demonstrates the viral genotypes can be utilized as molecular barcodes in combination with epidemiologic data to monitor the spreading routes of the pandemic and evaluate the effectiveness of control measures. Moreover, the dynamics of viral mutational spectrum in the study may help the early identification of new strains in patients to reduce further spread of infection, guide the development of molecular diagnosis and vaccines against COVID-19, and help assess their accuracy and efficacy in real world at real time.

## Introduction

A severe respiratory disease, named as Coronavirus Disease 2019 (COVID-19) has swept the globe by causing over 190 million confirmed cases (∼3.2% world population) and over 4 million deaths since it was firstly reported from Wuhan, China in early December 2019 [1]. A novel betacoronavirus, severe acute respiratory syndrome coronavirus-2 (SARS-CoV-2), was identified as the etiologic agent of COVID-19 [2,3]. Genomic sequencing results indicate that the genome of SARS-CoV-2 is closely related to two bat-derived SARS-like coronaviruses, RaTG13 (with 96.2% identity) and RmYN02 (with 93.3% identity), respectively collected in 2013 and 2019 in Yunnan province, southwest China [4,5]. Phylogenetic analysis also demonstrates that SARS-CoV-2 is similar to the two bat-derived SARS-like coronaviruses but distinct from SARS-CoV and MERS-CoV [3,5]. Thus, it is speculated that bats might be the original host of SARS-CoV-2, and other non-bat mammals such as pangolins might have been the intermediate reservoir [6]. Moreover, the first reported patient cluster of COVID-19 was epidemiologically linked to a seafood wholesale market in Wuhan, China, so the market has been assumed as the origin of the outbreak by representing an intermediate reservoir of SARS-CoV-2 [7]. However, epidemiological evidence doubted the market was the birthplace of SARS-CoV-2 [8,9]. During the 2014-2015 Ebola outbreak, full-length EBOV genome sequences from different severely stricken countries/districts in West Africa have helped us to better understand the viral evolution and transmission dynamics of the outbreak [10–12]. Likewise, genomic studies of SARS-CoV-2 viral sequences may provide key insights into the transmission and evolution dynamics of the ongoing COVID-19 pandemic.

In this study, we firstly analyzed 4,013 full-length genome sequences of SARS-CoV-2 submitted to the GISAID EpiFlu ™ database from all over the world (N = 4,002) and from NGDC database (N = 11) in China over a 14-week timespan since the outbreak in Wuhan, China (as of 7 April 2020). By mutation-based genotype characterization, we gain insights into the global evolutionary dynamics and genetic diversity of SARS-CoV-2 from the early phase of the pandemic. Moreover, we also used the same approach to analyze 261,323 full-length SARS-CoV-2 genomes from all over the world over 12 months since the outbreak (i.e. including all the available viral genomes in the database as of 25 December 2020) to recapitulate those insights in parallel with the unfolding pandemic. So this study not only provides an unprecedented window into the global transmission trajectory of SARS-CoV-2 in the early phase, but also reveals the subsequent expansion patterns of the pandemic.

## Materials and methods

### 1. Genome sequence retrieval and cleaning

We retrieved 4,555 FASTA sequences of SARS-CoV-2 genomes from Global Initiative on Sharing All Influenza Data (GISAID) database (https://www.gisaid.org/) and 147 FASTA viral sequences from National Genomics Data Center (NGDC) database (https://bigd.big.ac.cn/ncov) in China as of 7 April 2020, the first cutoff point of this study. The first sequenced viral genome Wuhan-Hu-1 (Genbank ID: MN908947.3) comprising 29,903 nucleotides with annotation of corresponding CDS regions was downloaded from Genbank (https://www.ncbi.nlm.nih.gov/genbank) as well as the related coronavirus genome sequences from bats and pangolins.

Partial SARS-CoV-2 genome sequences or gene-level only sequences were filtered out. Viral genome sequences from non-human hosts were also filtered out. Redundant sequences included in both databases, multiple samplings from the same patient, and re-submission of the identical sequences were excluded. Sequences with N for more than 3% of the total nucleotides (except 5’ and 3’ ends) were filtered out. After filtering, all the remaining sequences were mapped to the reference genome by a dual alignment software MAFFT (v7.450) which takes into consideration of both amino acid and nucleotide sequences. The genome sequences with >20 mismatches to the reference genome were further filtered out. After the filtering process, a total of 4,013 viral genome sequences (4,002 from GISAID and 11 from NGDC) were included in this study for further analysis.

Similarly, we retrieved 290,005 FASTA sequences of SARS-CoV-2 genomes from GISAID database as of 25 December 2020, the second cutoff point of this study. After the aforementioned filtering process (except the genome sequences with > 45 mismatches to the reference genome were filtered out here), a total of 261,323 viral genome sequences were included in this study for further analysis.

### 2. Phylogenetic tree analysis

In order to find evolutionarily related coronavirus with SARS-CoV-2, the first sequenced viral genome sequence (Genbank ID: MN908947.3) was used to perform BLAST via NCBI betacoronavirus sequence dataset (https://blast.ncbi.nlm.nih.gov/Blast.cgi). Nine coronavirus sequences from bats sharing the highest genomic identity with MN908947.3 were selected and downloaded. In addition, nine coronavirus sequences from pangolins (available at GISAID) were also selected to align with MN908947.3. Phylogenetic tree analysis was conducted with the neighbor-joining method in MEGA-X (v10.0) based on the alignment results. Six sequences (three from bats and three from pangolins) that are most proximate to MN908947.3 in the tree were chosen for nucleotide alignment at the orthologous sites of 8782 base and 28144 base with the sequences of nine early Wuhan cases (eight linked with the Market and one not related to the Market) [7].

### 3. Mutation calling and clustering analysis

Mutations were detected by an in-house developed software, which comparing each sample’s alignment result to the reference genome sequence. The first 150 base at 5’ end and 80 base at the 3’ end were omitted, and the ambiguous bases were ignored. After mutation detection, the matrix of mutations for all samples was used to perform the unsupervised clustering analysis via Pheatmap (v1.0.12) package of R.

### 4. Strain of Origin (SOO) algorithm

19 genotypes were selected from clustering analysis and defined in the Pedigree chart (**Fig. 4b**). Samples with mutation profiles matching to any of the 19 defined genotypes were classified into the corresponding types, whereas samples with mutation profiles not fitting into any defined genotypes were assigned as Others. Samples with no mutations under the aforementioned mutation calling methods were defined as ancestral type.

## Results

### 1. A super-dominant genotype of SARS-CoV-2 was characterized with two concurrent mutations in the early phase of COVID-19

By the first cutoff date (7 April 2020), we identified two most abundant substitutions, C/T at location 8782 base (orf1ab: C8517T, synonymous) and T/C at location 28144 base (ORF8: T251C, L84S) from 4013 viral genomes. The T8782 and C28144 genotypes were found to co-exist in 767 (19.1%) genomes, whereas the remaining 3,246 genomes (80.9%) were consistent with the first sequenced SARS-CoV-2 genome, Wuhan-Hu-1 (MN908947.3) at those two sites [2].

Next, to address the question of whether those two sites are evolutionarily conserved, we generated a phylogenetic tree of the eight patient samples linked with the Huanan Seafood Wholesale Market (hereinafter named as the Market) and the related coronaviruses from animal reservoirs by nucleotide sequence alignment [7,13]. Interestingly, we found the most related coronaviruses from pangolins and bats showed consensus at the orthologous sites of 8782 base as T and 28144 base as C. A complete linkage at both sites was also observed in these highly related coronaviruses including the most closely related bat coronavirus RaTG13 (96.2% identical) (**Fig. 1**). This result suggests that the T8782 and C28144 genotype existing in 19.1% of SARS-CoV-2 genomes is more conserved during evolution as an ancestral genotype. On the opposite, the samples from the eight patients demonstrate identical concurrent mutations on those two sites (T8782C and C28144T). Coincidentally, all eight patients had worked or visited the Market before the onset of illness. Also worth mentioning is that the patient of sample Wuhan/WH04/2020 did not visit the market but stayed in a hotel nearby between 23 and 27 December, 2019 [2,7]. Different from the aforementioned eight Market samples, the genotype of this patient sample showed no mutations on the two sites (i.e. T8782 and C28144), suggesting this patient had been infected from somewhere else in Wuhan instead of the Market. Noteworthily, the first sequenced SARS-CoV-2 genome, Wuhan-Hu-1 which was from a worker at the Market, also acquired the two point mutations [2]. Based on the analysis above, the most recent common ancestor (MRCA) of the SARS-CoV-2 should be Wuhan/WH04/2020 instead of Wuhan-Hu-1, although it was first sequenced [14]. Therefore, Wuhan/WH04/2020 was used as the reference genome for all subsequent analyses in the study.

**Fig. 1.**
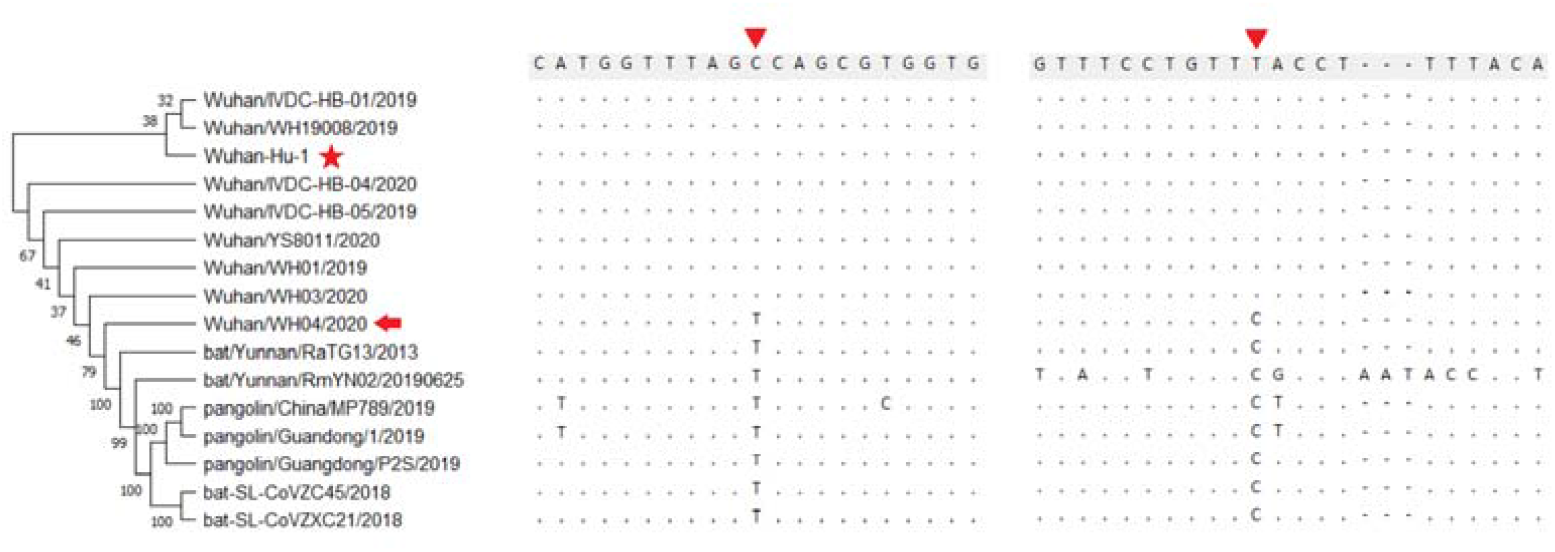
Sequence alignment of SARS-CoV-2 and the most related coronaviruses from animal reservoirs at location 8782 and 28144. The 20 base frank sequences of site 8782 and 28144 (indicated with red triangle) from eight COVID-19 samples linked with the Market, one sample not directly linked with the Market (Wuhan/WH04/2020) (marked with red arrow), and seven closely related virus samples from bats and pangolins were aligned with Wuhan/WH04/2020. The first sequenced genome, Wuhan-Hu-1 was marked with red star. Bat/Yunnan/RaTG13/2013 is the closest coronavirus to SARS-CoV-2 with overall 3.8% genomic difference.

Given limited sampling of viruses from the Market, we acknowledge that samples with concurrent T8782 and C28144 genotype from the Market might have been underrepresented. However, we are confident that a significant portion of samples from the Market were derived from an ancestral genotype, generating a distinctive genotype defined by two concurrent mutations, which we named as M type (T8782C/C28144T) hereinafter. It represented an overwhelming majority of all COVID-19 samples since the initial phase of the global pandemic (**Fig. S1**). All the 16 samples collected prior to 01 January, 2020 have the M type mutations that coincides with the fact that market contact history was one of the diagnostic criteria of COVID-19 at the period of time (**Table S1**) [9].

### 2. Viral genotypic composition of early cases reported from Wuhan were already diversified

Early cases reported from Wuhan was extremely critical to answer how the outbreak took place at the very beginning. In this study, we were able to collate 34 viral genomes sampled from Wuhan between December 24, 2019 to January 18, 2020, although the number of confirmed case by then were 121 according to Chinese officials (**Fig. 2**). There were two distinct clusters of the 34 early samples. 30 out of 34 viral genomes were categorized into the M type (T8782C/C28144T) with a great extent of genetic diversity. Among these 30 genomes, 17 acquired extra mutations apart from two M type mutations resulting in 14 different genotypes. All of the 11 viral genomes linked with the Market (including 8 samples of patients who had visited the Market and 3 positive environmental samples collected from the Market) were in this cluster [7]. Although the M type was the dominant type during the early outbreak of COVID-19 in Wuhan, the non-Market genotypes from four patients forms the second cluster that also co-existed with M type cluster at that time. Two of them were ancestral type and the other two had their own unique mutations. Wuhan/WH04/2020 was a patient who had no direct Market exposure in the second cluster [7]. Taken together, these findings imply that the genetic pool of SARS-CoV-2 was already very diversified during the early outbreak in Wuhan as there were 18 different genotypes in total among the 34 early samples from Wuhan. The super-dominant Market lineage might have been initially transmitted to the market by a primary patient case infected with the M type virus. M type virus was rapidly propagated within the Market which had served as a big incubator of the outbreak considering its huge size (∼50,000 square meters and ∼1000 booths). This notion is also evidenced by the three positive environmental samples (Wuhan/IVDC-HB-envF13/2020, Wuhan/IVDC-HB-envF13-20/2020, and Wuhan/IVDC-HB-envF13-21/2020) collected from the booths and garbage truck of the Market in 1 January 2020 by China CDC. The viral genotypes of three environmental samples were also M type. In fact, 33 out of 585 environmental samples from the Market were tested positive for SARS-CoV-2 according to an investigation conducted by China CDC in January 2020.

**Fig. 2.**
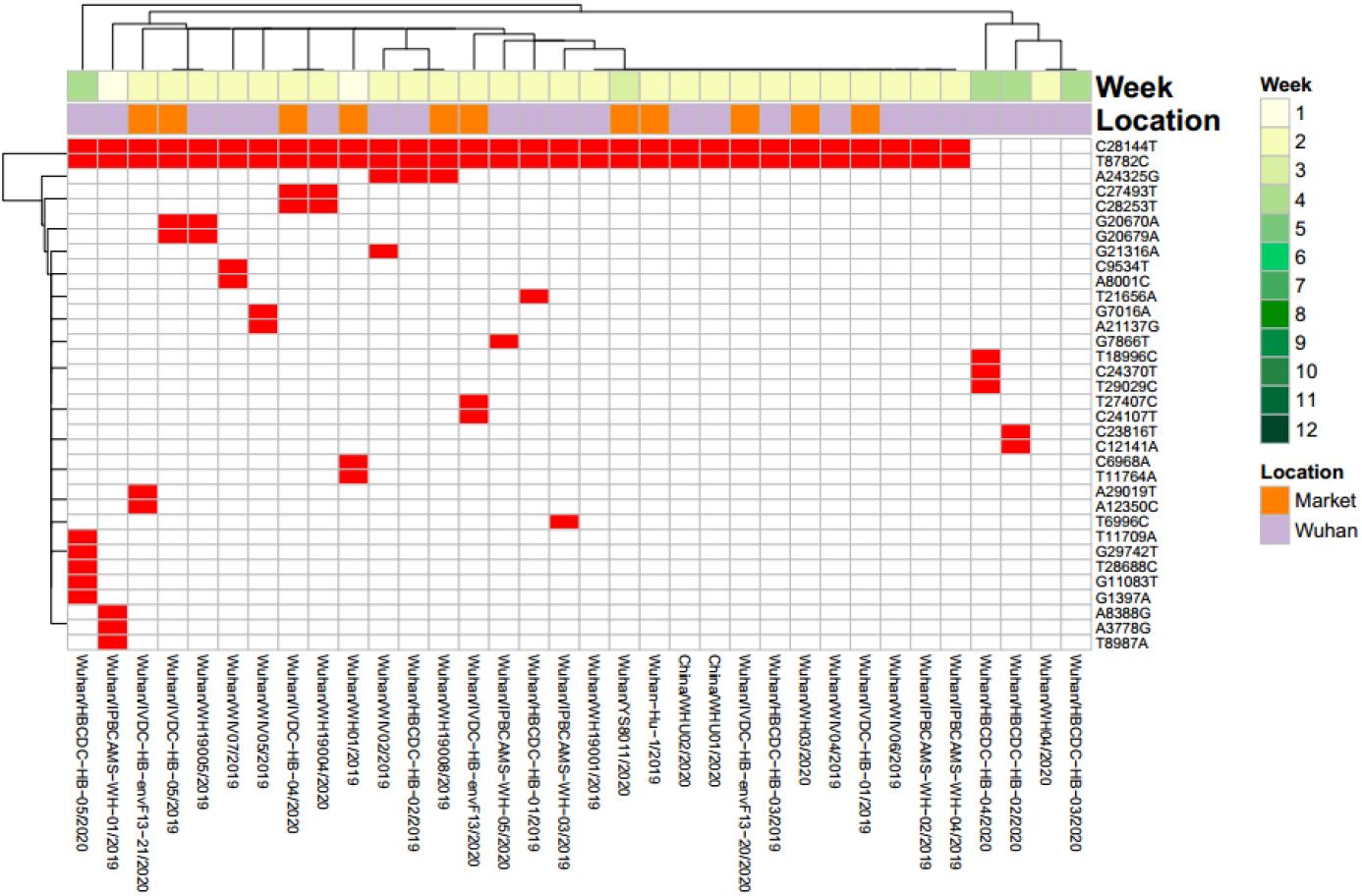
Mutation profiles of 34 early samples from Wuhan. Row one: time of sample emergence calculated as week(s) since the week of 23 December 2019; Row two: Sample location. Samples linked with the Market were indicated as orange.

### 3. Potential bias introduced by sequencing errors on mutation analysis was insignificant

Before jumping to any conclusive summary of the mutation spectrum of SARS-CoV-2 genome, the potential impact of sequencing errors on mutation calling in this study should be cautiously addressed. To estimate the magnitude of bias that may be introduced by sequencing errors, we firstly filtered out low quality sequence data to ensure the high quality of the analyzed genome sequences. The samples with only one of the concurrent mutations of M type at either position 8782 base or 28144 base were regarded as the consequence of sequencing/assembling errors since the co-mutations on the two sites were interrupted. Thus these samples were used to exemplify the maximum sequencing errors that can be anticipated. 16 out of 4,013 genomes bore only one of the two concurrent M type mutations, and six of the 16 demonstrated an ambiguous base (N or “Y”) at the inconsistent site. So there were supposed to be at most 10 genomes unambiguously bearing the correspondence of sequencing errors at either of the two sites in the 4,013 genomes. For simplicity we assumed sequencing errors occurred randomly along the viral genome, so that the maximum sequencing error rate for each base per genome can be calculated as 10/2/4013 = 0.00125. And the error rate was further divided by 3 given that each base was equiprobably recognized as one of the three erroneous bases (e.g. A->C, A->T, A->G), resulting in a final error rate of 0.00042. Based on this estimation, we can assume if any single mutation observed in this study had been caused by sequencing error, it was supposed to be found in no more than 1.68 genomes (calculated as 4013 genomes multiplied by 0.00042). Similarly, amongst 4013 genomes in the study, less than 0.001 genome was anticipated to erroneously acquire two concurrent mutations such as M type by sequencing errors (calculated as 4013 genomes multiplied by 0.00042^2).

As the evolutionary rate of SARS-CoV-2 was estimated to be 27.1 subs per year (see more details in the next section of results), it may take 3321 years (30000 × 3 possible alternative bases /27.1 bases per year) for a viral genome to generate one identical mutation with another one solely through viral error-prone replication process. Thus, less than one (0.4) out of 4,013 viral genomes was anticipated to acquire one identical mutation with another one by random mutation events during the past 4 months (calculated as 4013 genomes divided by 3321 years and multiplied by 1/3 year). In addition, as indicated previously, amongst the 4013 viral genomes, less than 0.001 genome was anticipated to acquire two concurrent mutations as M type by sequencing errors. It reinforced the notion that identical genotypes of any multitude observed in the study was very unlikely to have been caused by coincidence but resulted from lateral human-to-human transmission.

### 4. The mutation spectrum and dynamics of SARS-CoV-2 genome in the early phase of COVID-19

As described above, sequencing errors were very unlikely to confound the mutation analyses of 4,013 genomes. A total of 2,954 unique nucleotide substitutions were identified from the 4,013 SARS-CoV-2 genomes with relatively even distribution across the viral genome (**Table S1**). On average, there are 7.4±3.4 (mean±SD) mutations per genome. Only 31 genomes had no mutation (i.e. ancestral type), while 952 (32.2%) mutations were recurrent in more than one sample. There were 17 mutations that occurred in more than 10% samples (**Fig. S2, Table S1**).

Mutations increased in individual samples during the course of evolution by plotting the number of mutations per sample with the time of sample emergence (**Fig. S3**). Samples with more mutations were collected at a relatively later stage. A simple linear regression of the root-to-tip genetic distances against the sampling dates was performed to estimate the evolutionary rate of SARS-CoV-2 using the TempEst (v1.5.3) software. The evolutionary rate was estimated to be 27.1 subs per year, which was very similar to the evolutionary rate (26.7 subs per year) estimated by Nextstrain.org from 4616 viral genomes sampled between December 2019 and April 2020 (https://nextstrain.org/ncov/global?l=clock).

Like M type T8782C/C28144T mutations, concurrent mutations were also observed from the other 15 most frequent single nucleotide mutations. A symmetric matrix plot by clustering analysis was generated from the 17 most frequent mutations to highlight the most common concurrent mutations (**Fig. 3a**). T8782C/C28144T were concurrent in 81% samples, followed by C14408T/A23403G/C3037T/C241T (51%), G28881A/G28882A/G28883C (16%), C1059T/G25563T (12%), C17747T/A17858G/C18060T (12%) and G11083T/C14805T/G26144T (8%). G28881A/ G28882A/ G28883C and C1059T/G25563T were intersecting with C14408T/C241T/C3037T/A23403G since both were subsequent mutations of C14408T/C241T/C3037T/A23403G (**Fig. 3b**). Likewise, C14408T/C241T/C3037T/A23403G and G11083T/C14805T/G26144T were intersecting with T8782C/C28144T since both were subsequent mutations of T8782C/C28144T. C17747T/A17858G/C18060T didn’t intersect with any other concurrent mutations since it was a genotype derived directly from ancestral type and independent to T8782C/C28144T (M type).

**Fig. 3.**
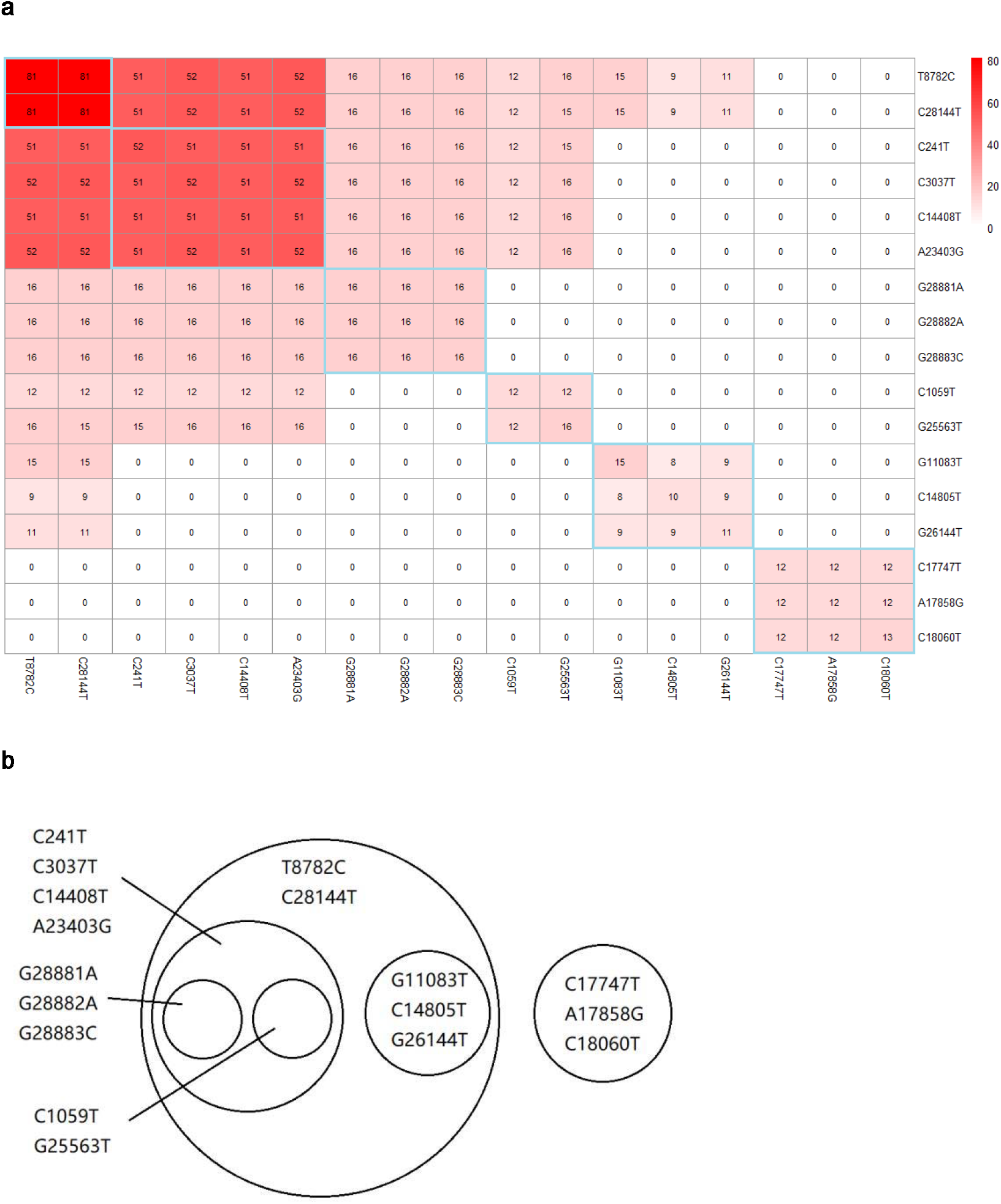
Concurrence and relationships of the 17 most frequent mutations. (a) Symmetric matrix of concurrence rate of 17 most frequent mutations among all samples. The number within each box represented the percentage of samples possessing the intersecting mutations against all samples. (b) A Venn diagram showing the subsequence relationships of the 17 most frequent mutations.

**Fig. 4.**
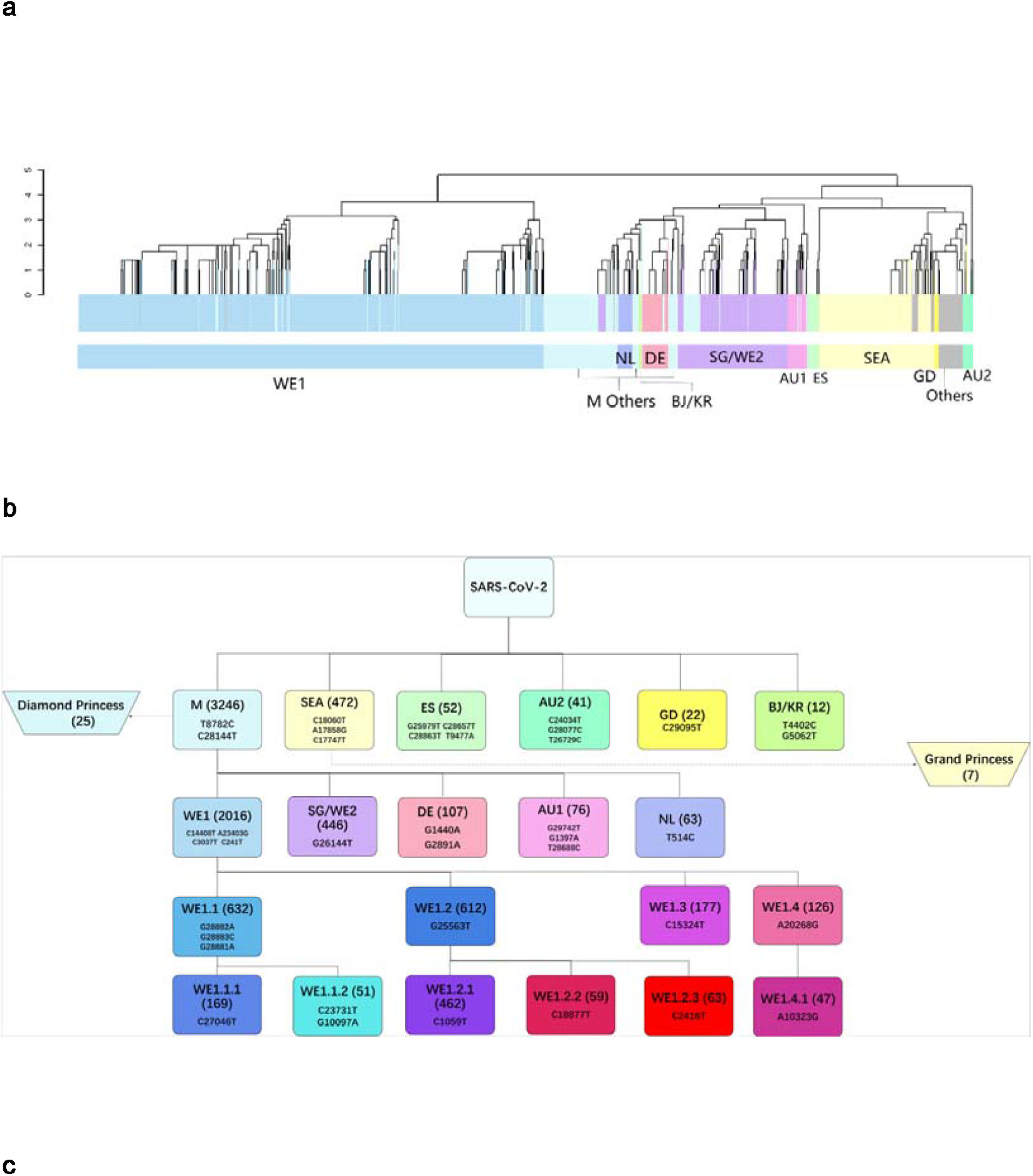
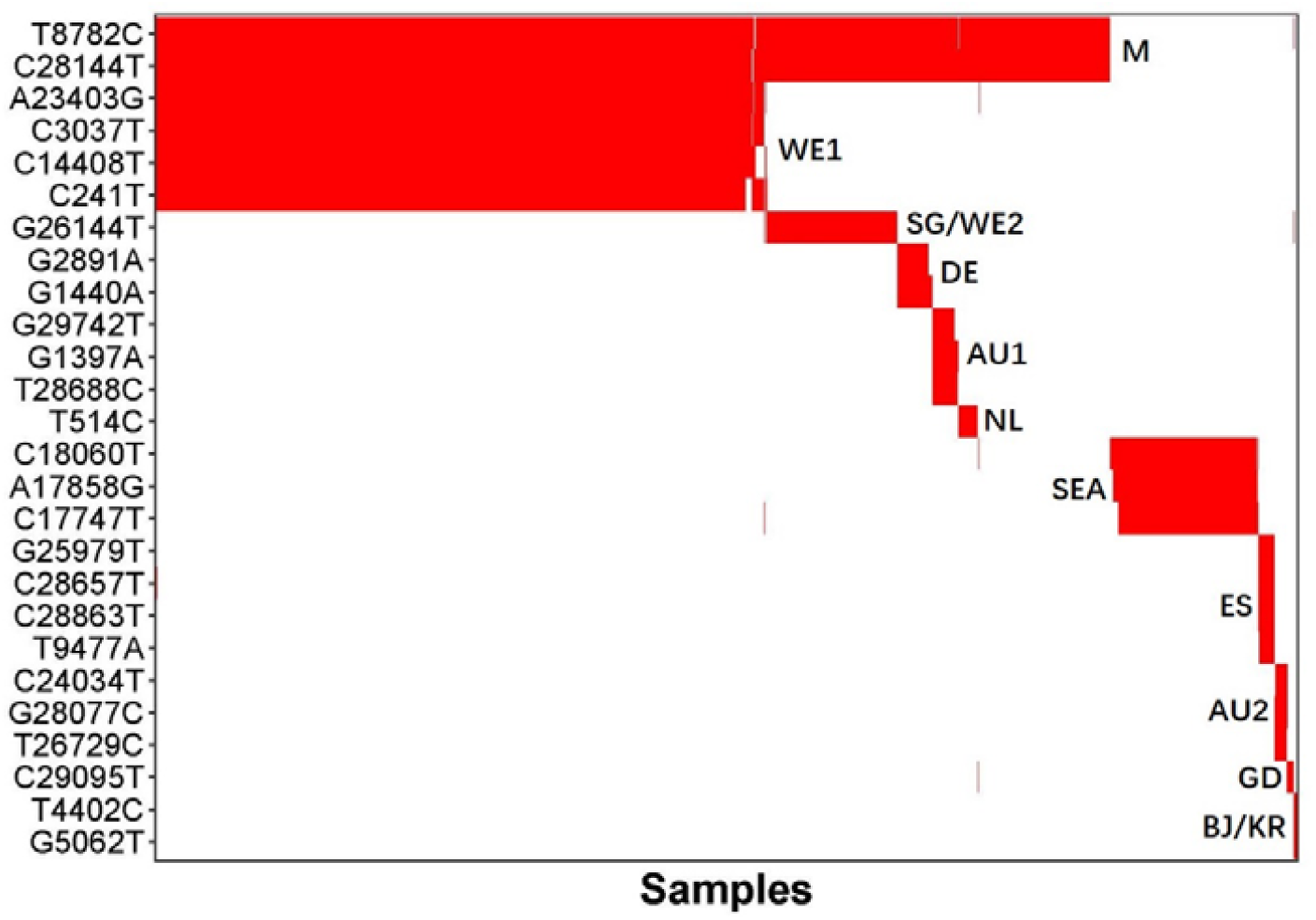
Major genotypes identified in the study. (a) Unsupervised mutaion clustering of all samples. Mutations concurrently called from at least 5 samples were included. 11 distinctive major mutation profiles were identified based on clustering tree branches and were named mainly based on the geographic locations where a certain genotype was initially or mainly reported from. The two-letter ISO country codes were used to indicate the countries associated with the mutation profiles (as shown at lower color bar). The upper color bar demonstrates genotypic homogeneity within each clustering tree branch. (b) Pedigree chart of major genotypes. In combination of mutation clustering and available epidemiologic information, 11 distinctive main genotypes were characterized and the pedigree chart demonstrated the relationship of each genotype. The genotypes from Diamond Princess and Grand Princess derived from M type and SEA type respectively, were indicated with dashed arrows. (c) Mutual exclusivity among the major genotypes of Level-1 (M, SEA, ES, AU2, GD and BJ/KR) and Level-2 (WE1, SG/WE2, DE, AU1 and NL). M-Others: the other minor genotypes in Level-2. See also Fig. S4.

#### 4.1 The super spreading genotypes

952 mutations (32.2%) spread at least once as they were detected in more than one patient samples, and distinct genotypes can be characterized based on the prevalence of mutations (**Table S1**) in order to identify “super spreaders” with particular genotypes, who to a great extent determined the scale and trend of the ongoing pandemic. Super spreader genotype was methodically defined as the basal outbreak variant possessing certain common mutations, which caused the outbreak with a single introduction and subsequently transmission and evolution.

The biggest super spreader genotype was the variant carrying M type (T8782C/C28144T) mutations which was spread into 3,246 patients, counting for 80.9% of the cases in the study (**Fig. S1**). It may worth pondering whether this “founder effect” was attributed to a single super spreader or a multitude of coincidentally identical super spreaders. Based on our aforementioned estimation, the chance to have a multitude of identical mutations by coincidence or sequencing error was next to none. Therefore, it is plausible to assume that the patient clusters from the Market in Wuhan during the early outbreak were very likely to be the descendants from literally one single ancestor, who might have been a vendor or a regular customer and probably spread the virus at the Market late November or early December according to the limited epidemiologic data.

In contrast, only 41 patients (1.0%) had unique genotypes that were not overlapping with any other mutations identified in the 4,013 viral genomes (i.e. singletons) (**Table S1**). These patients had not spread their viruses based on the sampled viral genomes in the early phase of the pandemic.

#### 4.2 Mutation-based unsupervised clustering identified major genotypes of SARS-CoV-2

In order to trace the temporospatial transmission and regional expansion the pandemic, we conducted mutation-based unsupervised clustering of all the samples. As shown in **Fig. 4a**, the 4,013 samples were largely grouped by their mutation profiles. Based on the clustering result, a pedigree chart of five-level hierarchy was manually curated and illustrated to show how the most dominant descendent genotypes were derived from the ancestral genome of SARS-CoV-2 at the Level-0 of the hierarchy (**Fig. 4b, Fig. S4**). Six descendant genotypes, namely M type (concurrent T8782C/C28144T) starting from the Market, SEA type (concurrent C17747T/A17858G/C18060T) initially reported solely from the Greater Seattle area in USA, ES type (concurrent T9477A/G25979T/C28657T/C28863T) with more than 60% of cases reported from Spain, AU2 type (concurrent C24034T/T26729C/G28077C) with 22 out of 41 cases reported from Australia, GD type (C29095T) initially found in Guangdong Province, and BJ/KR type (12 cases with concurrent T4402C/G5062T mutations) reported from both Beijing and South Korea, were directly derived from the ancestral genome by acquiring corresponding mutations, and thus represented Level-1 of the hierarchy (**supplementary results**).

Five descendant genotypes (WE1, SG/WE2, DE, NL, AU1) were further derived from M type, consisting of Level-2 of the hierarchy. In particular, the most prevalent descendant genotype of M, WE1 type (named as Western European 1), represented a total of 2,016 cases, a half of all the cases (50.2%) in the study. Over 70% of WE1 cases were reported from Western European countries, with the United Kingdom (19.2%), Iceland (11.8%), Belgium (9.7%), France (8.5%), and Netherlands (5.0%) being the most severely stricken countries by WE1. The WE1 type was featured by four concurrent mutations (C241T/C3037T/C14408T/A23403G). Given geographic proximity among those countries, cross-border virus traffic might have occurred, leading to widespread transmission of SARS-CoV-2 in Western Europe. WE1 also represented 34.8% of the cases in the United States. Interestingly, among 4,013 samples, we found three early samples carrying three out of the four mutations of WE1 (C241T/C3037T/A23403G), with two (one from Germany and one from Shanghai) sampled on 28 Jan 2020 and one from Shanghai sampled on 31 Jan 2020. The one from Germany belonged to the first COVID-19 cluster reported from Bavaria, Germany, which was associated with a primary case with previous travel history from Wuhan [15].

SG/WE2 type was characterized by a single common mutation (G26144T) and was first reported from Singapore and several Western European countries (UK, France, Switzerland and Netherlands). All early cases of the DE type (featured with concurrent G1440A/G2891A) were found in Germany, however, the majority (62.6%) of DE cases were reported from UK. NL type was mainly reported from Netherlands and featured with a single extra mutation, T514C. AU1 type was mainly found in Australia with three extra concurrent mutations (G1397A/T28688C/G29742T).

**Fig. S5** illustrated the temporal expansion of the 11 major genotypes over 14-week timespan. M type remained as the overwhelmingly dominant genotype from the very beginning of the outbreak to early April. WE1 was spread to more than half of the total cases as of 7 April, becoming the most prevailing M-derived genotype in the globe. Next to WE1 type, SG/WE2 type was spread to 11.1% of global population. The major non-M type, SEA type, initially reported from the Greater Seattle area, was spread to 11.8% of global population. Moreover, genotypic compositions of SARS-CoV-2 in different countries and geographic locations were able to indicate genotypic-epidemiologic relevance (**supplementary results, Fig. S6-S8**).

Interestingly, as shown in **Fig. 4c**, six Level-1 genotypes (M, SEA, ES, AU2, GD and BJ/KR) derived directly from ancestral type were mutually exclusive, and five Level-2 genotypes derived from M type (WE1, SG/WE2, DE, AU1 and NL) were mutually exclusive as well. It implies mutations occurred randomly and independently in the genome of SARS-CoV-2 and the various genotypes carrying specific mutations were propagated during human-to-human transmission, not by accumulating hot-spot mutations during the replication of individual viral genomes. This also reflects the high quality of sequencing data applied in the study (after filtering out low quality sequence data) and the randomness of the mutations occurring across the viral genome.

#### 4.3 Genotype matching and strain of origin

By taking into account all of the well-defined 11 major genotypes from Level-1 and 2 in the study, we developed an algorithm, Strain of Origin (SOO), to match a particular SARS-CoV-2 viral genome to the known genotypes based on its mutation profile. The concordance of SOO was estimated in comparison with mutation clustering by assigning each of 4,013 samples included in the study to the corresponding genotype (**Fig. 5**). The overall concordance of genotypes assigned by SOO with those assigned by mutation clustering was 89.8%. Within Level-1 genotypes, the concordance ranged from 84.9% to 100.0% with an overall concordance of 86.5%. All the Level-2 genotypes represented major subtypes of M type and the overall concordance with clustering results at this level was 90.5%. The most abundant genotype at Level-2, WE1 showed 93.4% concordance. Thus, SOO represents a more accurate approach to define genotypes as it only takes into consideration the specific mutations of the particular genotypes with little influence from the rest random mutations.

**Fig. 5.**
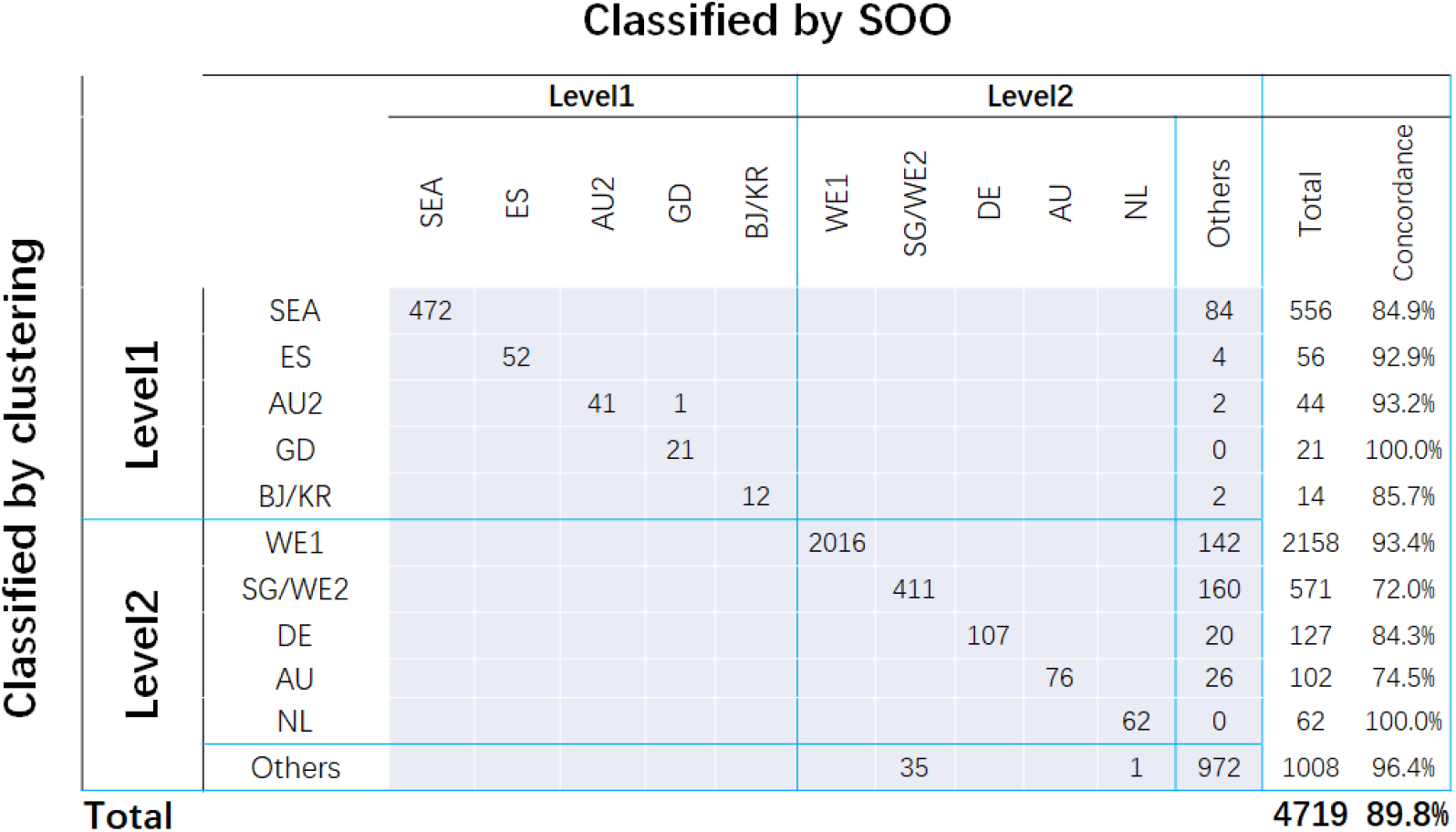
Performance of SOO algorithm in genotype classification. Agreement of genotype assignment between the clustering (y-axis) and SOO (x-axis). Percent of correct assignment from the total samples for each genotype from clustering was indicated to the right of the plot.

#### 4.4 Comparison of SOO classification and GISAID phylogenetic clades

There were seven phylogenetic clades of 5139 virus genomes sampled between December 2019 and September 2020 in GISAID global analysis (https://www.gisaid.org/epiflu-applications/hcov-19-genomic-epidemiology/) (**Table S2**). Since genomes were equally subsampled from each admin division per month, it should be acknowledged that countries with massive viral genome submissions might have been underrepresented. Thus conclusions regarding the global vision of the pandemic based on the GISAID global analysis should be drawn with caution. Four clades (n=4077, 79.3%) from GISAID were well defined by the SOO classification (**Table S3**). The three most prevalent clades GR (n=1726, 33.6%), G (n=1252, 24.4%) and GH (n=977, 19.0%) were descendent from WE1 (n=3955, 77.0%), with GR referred to WE1.1, GH referred to WE1.2, and G referred to WE1 others by SOO classification. Moreover, V (n=122, 2.4%) was referred to SG/WE2 by SOO. Two other clades can not be directly referred to any SOO genotypes although L can be vaguely inferred as ancestral type and others, S be inferred as a mixture of non-M types including SEA, ES, AU2, and GD. On the other hand, the O (n=500, 9.7%) clade can not be equivocally inferred as any SOO genotypes. It is plausible since it was not presented as a unique branch as other clades but scattered all over other branches of the phylogeny, implying it was not a well-defined unique clade.

### 5. The mutation spectrum of the subsequent one-year global expansion of COVID-19 pandemic

We analyzed all the available SARS-CoV-2 viral genomes in GISAID database as of 25 December 2020, the second cutoff date of the study. A total of 10,392 unique nucleotide substitutions were identified from the 261,323 SARS-CoV-2 genomes (**Table S4**), which indicates roughly one out of three nucleotides in the viral genomes has mutated during the 12-month timespan of the viral evolution. A pedigree chart of the 100 most abundant mutations was generated to highlight the lineages of the most common concurrent mutations during the 12-month time window of the unfolding pandemic (**Fig. 6, Table S5**). A very tiny proportion (92 genomes, less than 0.04%) of viral genomes were ancestral type. 59 (64.1%) of them were reported from China between January and March 2020, among which seven were sampled from Wuhan in January 2020 (**Table S6**). Despite of the overwhelming dominance of M type (95.3%), other major genotypes at Level-1 hierarchy in the early phase gradually faded out as the pandemic unfolds (**Fig. 3b, Fig. 6, Table S5**). For example, SEA type was one of the most common viral genotypes in early April, accounting for 11.8% of the total samples. However, the percentage of SEA type drastically dropped to only 1.0% by the end of December (**Fig. 6, Table S5**). Moreover, the proportion of other non-M mutations (ES, AU2, GD and BJ/KR) at Level-1 were too small to be listed within the 100 most common mutations (**Fig. 6**). Similarly, WE1 (88.6%) was still the major subtype of M (accounting for 93.0% of M) while other subtypes (SG/WE2, DE, AU1, and NL) at Level-2 gradually faded out. But still, four subtypes of WE1, namely WE1.1 (34.3%), WE1.2 (19.3%), WE1.3 (22.2%) and WE1.4 (6.1%) were reasonably represented. On the other hand, a subtype of WE1, named WE1.5 (19.9%) featured with additional seven concurrent mutations (T445C/C6286T/G21255C/C22227T/C26801G/C28932T/G29645T) had not emerged by early April but came to the surface during the subsequent expansion. Interestingly, one-time concurrence of more than four mutations like WE1.5 was seldomly represented in the early viral samples, but was more frequently observed in the later-phase samples (**Fig. 6**).

**Fig. 6.**
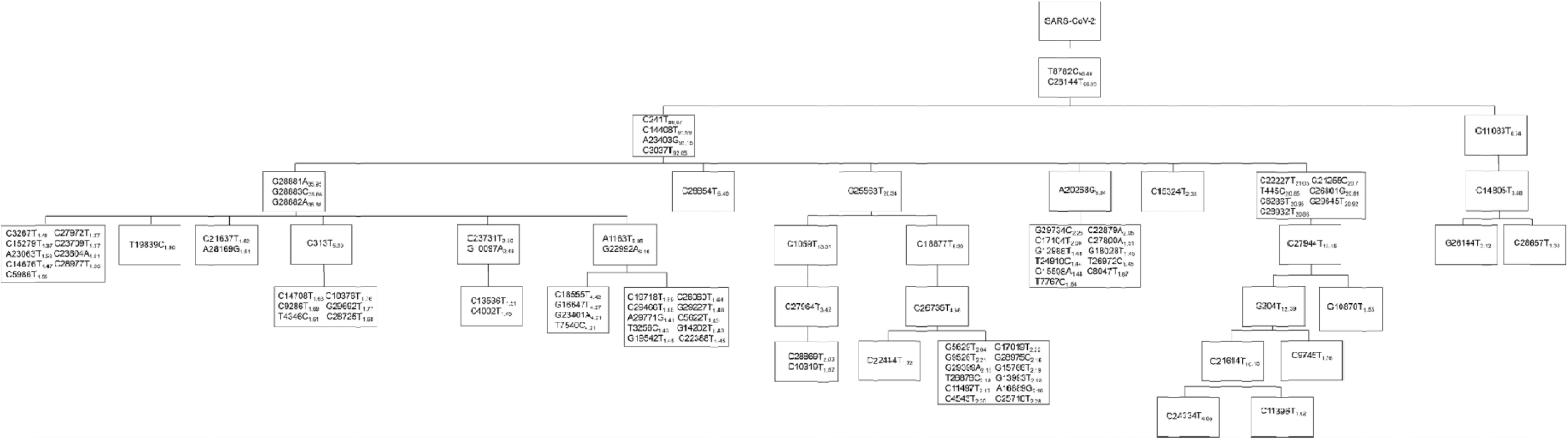
Pedigree chart of the 100 most abundant mutations from the 261,349 SARS-CoV-2 genomes (as of 25 December 2020). The number next to each mutation indicates the frequency of the mutation.

### 6. Lineage analysis of SARS-CoV-2 virus in the early phase sheds light on subsequent expansion of COVID-19 pandemic

In order to visualize the pandemic expansion patterns in the world over a significant one-year time window, we analyzed time-series genotypic compositions of SARS-CoV-2 at critical time points to try to piece the puzzle together (**Fig. 7, Table S1, Table S4**). First, to better fathom the whole story, a putative ‘patient zero’ harboring an ancestral viral genotype was added to build the first time point as 17 November 2019, on which date the earliest patient ever documented can be traced back to [16]. A total of 19 viral genomes were sampled by 1 January 2020, all of which were M type. As discussed before, M type cases had been populating at the Market for several weeks before it was shut down on 1 January 2020, resulting in an absolute overrepresentation of M type samples by this date. As the virus kept unfolding in Wuhan, the city was lock down on 23 January 2020. 80 of 104 (76.9%) viral samples by then were M type. Population mobility from Wuhan before its lockdown (most likely during the Spring Festival travel rush) may have caused the subsequent national-wide epidemic in China and ultimately the global pandemic. By 7 April 2020, over 80% of global cases were still M type. Noteworthily, a descendant genotype from M, WE1 type accounted for over a half of global cases. It firstly swept Western Europe in mid-February and later USA in late February, and became the most prevailing type worldwide. By 25 December 2020, 95.3% of global cases were M type and 93.0% of M-type cases were WE1, from which four VOCs were subsequently derived. The current overwhelming dominance of M (particularly one subtype WE1), and its continuous expansion were well captured and characterized throughout the study.

**Fig. 7.**
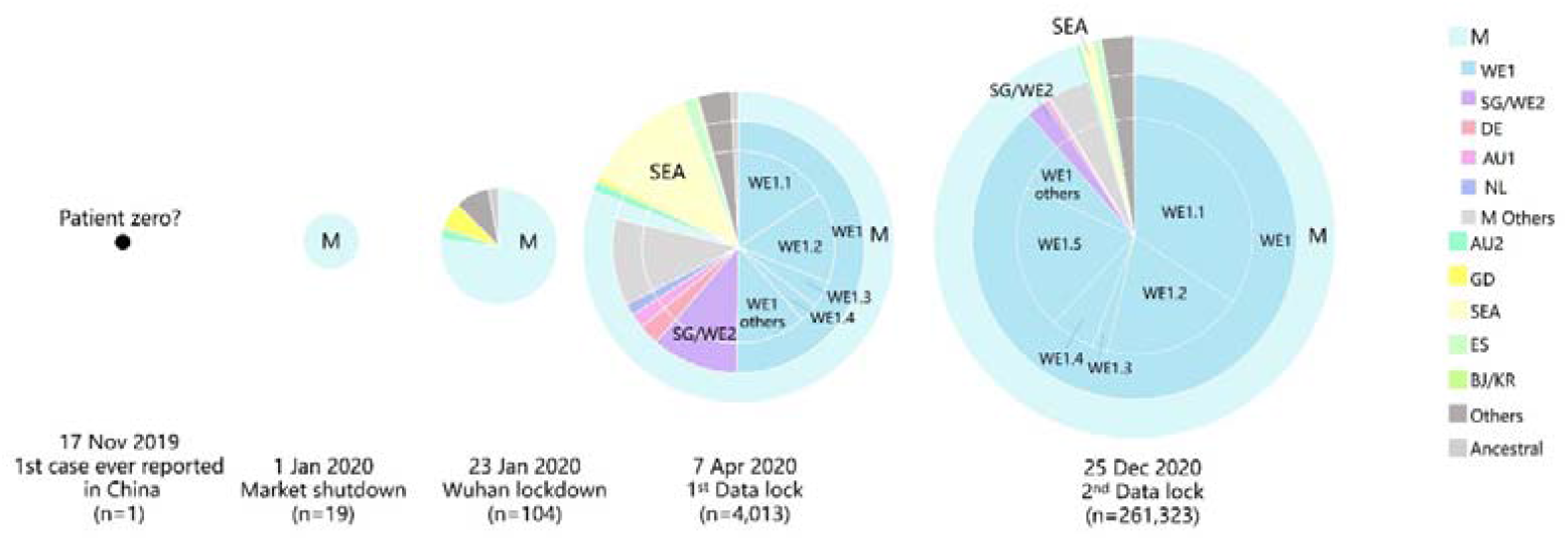
Snapshots of genotypic compositions of SARS-CoV-2 at five critical time points. Genotypic compositions of genomes sampled by the dates of the first case ever reported, market shutdown, Wuhan lockdown, the first data lock and the second data lock of this study were analysed.

Thus, the M type expansion pattern well represented the pandemic expansion pattern: patient zero (17 November 2019, unknown%) ➔ Market (1 January 2020, 100%) ➔ Wuhan (23 January 2020, 76.9%) ➔ World (7 April 2020, 80.9%; 25 December 2020, 95.3%).

### 7. The worrisome four Variant of Concerns (VOC) at present were all descendants from WE1

Recently, there have been four variants of concern (VOC) which draw tremendous public attention due to increased transmissibility or virulence that may attenuate effectiveness of current control measures, available diagnostics, vaccines, and therapeutics. In May 2021, the WHO has officially renamed the four VOCs based on the Greek alphabet for purposes of public discourse, in which UK variant N501Y V1 (i.e. B.1.1.7) was named Alpha, and the South Africa variant N501Y V2 (i.e. B.1.351) was named Beta. The two other VOCs were Gamma (i.e. P.1), the variant first identified in Brazil, and Delta (i.e. B.1.617.2) that originated in India [17]. In this study, a total of 4130 viral genomes harbored N501Y by 25 December 2020. Interestingly, they were generally categorized into two strains, with 3931 genomes as a subtype of WE1.1, and 188 genomes as a subtype of WE1.2 under SOO algorithm (**Fig. 8a**), with the former mainly reported from the UK (98.5%) and the latter mostly reported from South Africa (96.3%). It was consistent with a previous report from WHO that the UK variant N501Y V1 (i.e. B.1.1.7) was a different virus variant from the one from South Africa N501Y V2 (i.e. B.1.351) by phylogenetic analysis [18]. Interestingly the first genome of V1 in GISAID was from Victoria, Australia on 3 June 2020. A total of 31 V1 genomes were identified in June 2020, 30 of which were from Australia. This gave rise to the first wave of V1 in June. It followed by a huge spike beginning in November 2020 which was attributed to the wide spread of V1 in the UK at that time (**Fig. 8b**). The first genome of V2 was actually from New York City, USA on 21 April 2020, and V2 was later widely spread in South Africa as evidenced by a wave of V2 in November (**Fig. 8b**). In contrast, neither Gamma and Delta variants were found in the 261,323 viral genomes by 25 December 2020 in this study, which is plausible since they emerged later than Alpha and Beta variants.

**Fig. 8.**
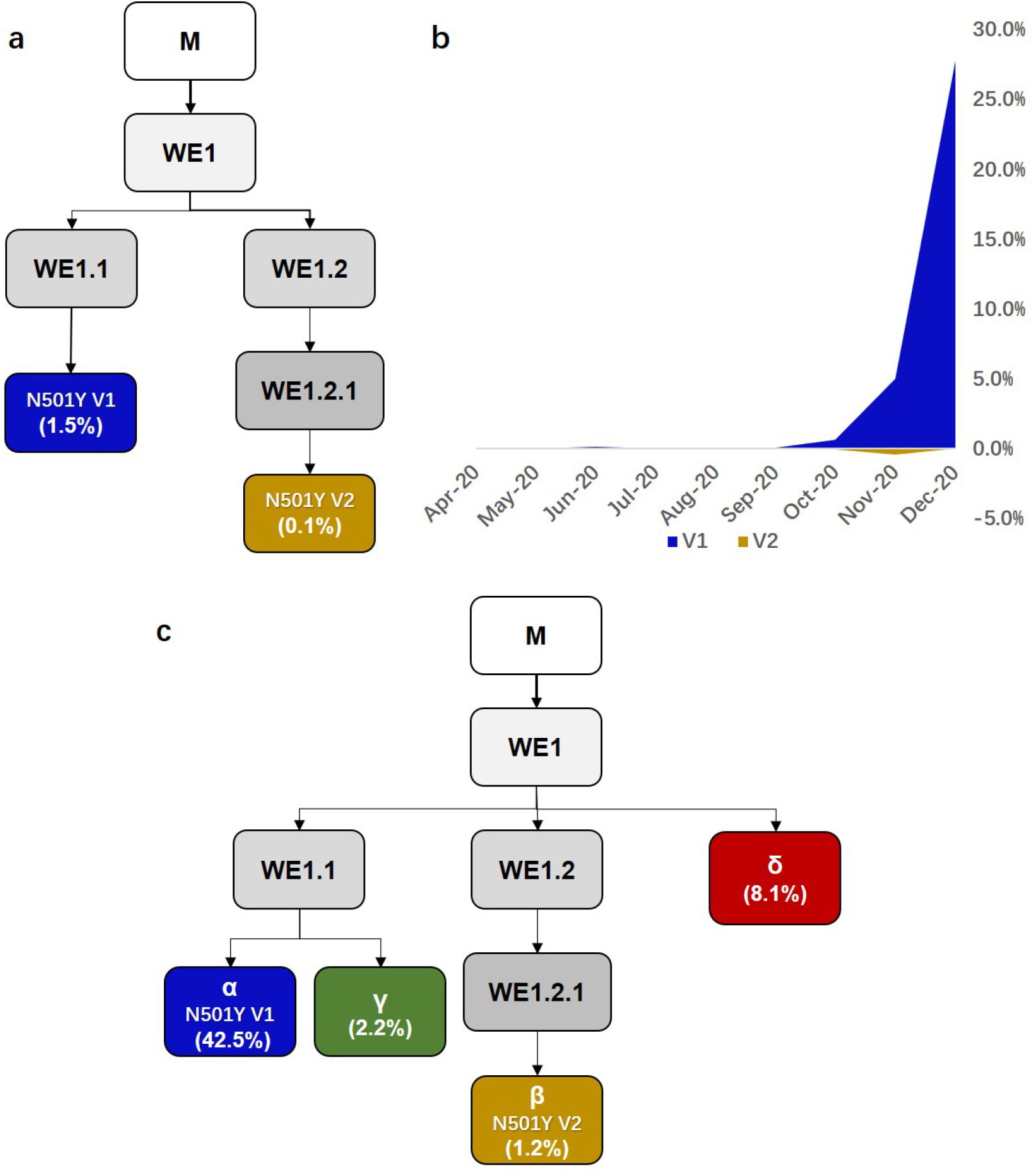
Emergence of VOCs. (a) The UK variant (N501Y V1) and South Africa variant (N501Y V2) were classified as two different strains by SOO. (b) The monthly frequency of V1 and V2 from April to December 2020. Percentages within the brackets indicate the number of each variant genomes against a total of 261,323 genomes as of 25 December 2020. (c) The four VOCs were classified by SOO. Percentages within the brackets indicate the number of each variant genomes against a total of 2,365,392 genomes as of 15 July 2021.

Next, to bridge the latest consequential situation of VOCs to what have been recapitulated in the one-year pandemic in the study, a *post hoc* analysis of the four VOCs under SOO algorithm was performed. We found that all the VOCs were subtypes of WE1 (**Fig. 8c**). In particular, the Alpha variant is the most prevailing VOCs (accounting for 42.5% of all the 2,365,392 genomes as of 15 July 2021) and is a subtype of WE1.1 in parallel with Gamma which is much less prevalent (2.2%). The Beta variant on the other hand, is a subtype of WE1.2.1, with the least prevalence (1.2%). The Delta variant was derived from WE1, a subtype not characterized by the SOO algorithm, which is understandable given that it just recently emerged. Although it accounts for a moderate amount (8.1%), it is still worrisome since it has been spreading faster than other variants [19, 20]. Moreover, featured with amino acid changes at the N-terminal domain (NTD) and the receptor binding domain (RBD) of spike protein, Delta variant has showed increased immune evasion potential [20].

## Discussion

It has been over seventeen months since Chinese health authorities first reported patient cases with pneumonia of unknown cause in Wuhan on 31 December 2019 [3]. As the pandemic continues, in order to mitigate the risk of further regional expansions as well as to estimate the effectiveness of control measures in various regions, researches on its origins, transmission routes and expansion models have begun to surge.

We acknowledge that the public available SARS-CoV-2 genomes included in this study are not sampled in strict proportion to the real-time global burden of COVID-19, however, we provide a global view of how the mutation patterns of SARS-CoV-2 genome vary over time in different countries and regions, which can shed light on the underlying temporospatial transmission and expansion pattern of COVID-19 worldwide. This is hitherto the largest and the most comprehensive SARS-CoV-2 viral genome study and molecular epidemiology study since COVID-19 outbreak in Wuhan, China. The 14-week timespan since the outbreak gives us a critical time window to study the mutation profiles and molecular evolution of SARS-CoV-2 at the initial stage of the pandemic.

SARS-CoV-2 virus is a positive-sense single-stranded RNA ((+)ssRNA) virus with a 30-kilobase genome, and like most other RNA viruses such as Ebola virus, SARS-CoV-2 can also quickly generate mutations through error-prone replication [2,3]. Considerable mutation events can be anticipated during the transmission and replication of the ongoing SARS-CoV-2 outbreak. Several studies on the genomics of SARS-CoV-2 virus have offered clues of the origins, and transmission path of the disease. However, due to lack of early samples, a limited number of SARS-CoV-2 genome and/or focusing on specific geographic locations [7,21–25], a complete global view of viral genomes in the context of their mutational spectrum is yet to be elucidated from any previous SARS-CoV-2 studies.

The very early cases of COVID-19, especially those linked to the Market, were the key to reveal the origin, the transmission paths as well as the evolution of the virus. Unfortunately, the viral samples and epidemiologic data from the early outbreak were largely mutilated. Here we meticulously collected the genomic data of the early cases from different databases and combed through the clinical data of those cases by not only in-depth review of early publications, but also collecting the viral sequences not included in GISAID, reading news reports and social media in China, and by contacting the researchers who worked on the early cases directly. We were able to collate 34 invaluable early cases from Wuhan including the cases of the patient cluster from the Seafood Market. The genetic diversity observed from the early Wuhan cases suggests the transmission had already been ongoing for some time at an inconspicuous pace before the clustered cases emerging from the Market were reported. The speculation can be reinforced by a report that the earliest patient can be traced back to 17 November 2019 [16].

Based on our time-series genotypic composition analysis, a super spreader genotype, M type, had ignited COVID-19 outbreak from the Market. The transmission continued for a few weeks or so without effective control measures until the final shutdown of the Market on January 1, 2020 [9]. As a consequence, the Market, with tens of thousands of people (workers and customers) in and out every day, became an incubator that catalyzed propagation of the M type in Wuhan during the early outbreak. Based on our lineage analysis, we can also conclude the explosion of the M type is the largest driving force for the global pandemic. However, it should be pointed out that this study is not equipped to address the natural origin of SARS-CoV-2 since there were no intermediate samples to link the most closely related bat coronavirus RaTG13 and human SARS-CoV-2.

The sequential increment of concurrent mutations from early lineages to descendent lineages as the pandemic unfolds still remains as an enigma. This phenomenon can be exemplified with M type. It initiated with 2 concurrent mutations followed by acquiring 4 concurrent mutations to become WE1, and further obtained 3 concurrent mutations to become WE1.1. Although it is roughly consistent with the estimated evolutionary rate (∼22 subs per year according to Nextstrain by December 2020), the underlying mechanism of those sequential increment of concurrent mutations is yet to be carefully unveiled.

Interestingly, we found that all four VOCs were subtype of WE1. Although the Alpha variant, a subtype of WE1.1, is currently the most prevailing VOCs, the Delta variant, a subtype of WE1, is more alarming since it spreads faster than other variant with increased immune evasion potential. Recent studies have demonstrated Delta variant spread is associated with an escape to antibodies targeting non-RBD and RBD Spike epitopes [19,20]. Yet more mechanistic studies are need to elucidate how it will affect the effectiveness of current control measures, diagnostics, vaccines, and therapeutics.

As many Emergency Use Authorized (EUA) real-time RT-PCR diagnostic tests for SARS-CoV-2 have been widely used all over the world to screen for infected COVID-19 patients, various genomic regions were chosen by different agencies and manufacturers to design primers for the tests. For example, the three target regions of the diagnostic kit developed by US CDC are within the N region, whereas the test that China CDC developed for the initial investigation in Wuhan targeted ORF1ab as well as the N region, which is similar to the test used in Singapore [3, 26]. On the other hand, many manufacturers’ tests chose to target the S gene. For example, the Thermo Fisher Scientific and Applied DNA Sciences tests target the S gene. Thermo’s test also targets the N and ORF1ab genes, while Applied DNA’s test targets two regions in the S gene [27]. Since genetic variants of SARS-CoV-2 arise regularly, those tests may give rise to potential false negative results due to the mutations in the viral genome. A few tests have been reported with false negative issues like S-gene dropout or reduced sensitivity with the S-gene target in detecting variants with N501Y mutation [27]. Not to mention tests detecting a single target in the viral genome which may generate far more variable and equivocal results. Although tests with multiple genetic targets to determine a final result are less likely to be impacted by increased prevalence of genetic variants, ongoing analyses of viral genomes in a real-time fashion may help with early identification of new stains in patients to reduce further spread of infection, guide the development and assess the efficacy of SARS-CoV-2 vaccines [28]. Based on our study, it is evident that the common mutation loci should be avoided as targets when designing RT and PCR primers for SARS-CoV-2 tests. Similarly, when developing nucleotide-based vaccines of SARS-CoV-2, researchers should take into consideration the mutation frequency in selecting viral genomic regions encoding antigen epitopes. Finally, it is imperative to reassure the vaccines can generate equivalent immunity against different genetic variants before inoculated in large population [19,20,28].

This study also demonstrates the genotypes of SARS-CoV-2 are unique identifiers that can be used as molecular barcodes to trace the virus transmission retrospectively and to reveal its expansion prospectively at the molecular level. The Strain of Origin (SOO) algorithm can match any particular SARS-CoV-2 viral genome to known genotypes with high accuracy based on its mutation profile. With the pandemic still ongoing, novel genotypes other than what we have characterized in this study may surface. Thus, we anticipate incorporating those newly emerging genotypes into the current algorithm may improve the performance of SOO in the future.

The United Kingdom launched a national SARS-CoV-2 Sequencing Consortium with £20M funding in March 2020, aiming to investigate how coronavirus is spreading in the UK, to help guide treatments in the future, and to anticipate the impact of mitigating measures [29]. The other countries have been following the same strategy. The results that we presented here serve as a proof of concept to demonstrate the utility of large-scale viral genome sequencing during a novel pathogen outbreak. Ramping up sampling in a real-time manner like the UK viral genome sequencing consortium may generate high-resolution maps of who-infected-whom transmission at community level and reveal the subsequent expansion patterns which are especially crucial for the most severely stricken countries and regions to promptly develop tailored mitigation plans [30].

## Author contributions

M.M. and Y.C. conceived and initiated the project. Under the guidance of M.M., S.L. and S.G. collected and analyzed the SARS-CoV-2 sequence data, and generated the figures and tables with the help from W.W. M.M. and Y.C. interpreted and refined the main results. Y.C. wrote the manuscript with critical review from M.M.

## Acknowledgements

We first would like to express our gratitude to the GISAID database and the NGDC database for collecting and sharing the SARS-CoV-2 sequence data. We also thank all the medical and research institutions from all over the world that promptly submitted the SARS-CoV-2 sequences to the aforementioned databases. We thank Dr. Shuyu Li for helpful comments on the manuscript.

## Data availability

The authors declare that all the SARS-Cov-2 FASTA sequence data utilized in the study are available in GISAID database (https://www.gisaid.org/) and Chinese NGDC database (https://bigd.big.ac.cn/ncov?from=groupmessage&isappinstalled=0). The other data supporting the findings of this study are available within the paper and its supplementary information files.

## Code availability

All the computer codes used in this study are listed as following, which are commercially available:

MAFFT v7.450 for sequence alignment;

Pheatmap (v1.0.12) R package for unsupervised cluster analysis;

MEGA-X v10.0 for phylogenetic analysis;

TempEst (v1.5.3) for evolutionary rate estimation.

## Additional resources

None

